# Layer-dependent activity in human prefrontal cortex during working memory

**DOI:** 10.1101/425249

**Authors:** Emily S. Finn, Laurentius Huber, David C. Jangraw, Peter A. Bandettini

## Abstract

Working memory involves a series of functions: encoding a stimulus, maintaining or manipulating its representation over a delay, and finally making a behavioral response. While working memory engages dorsolateral prefrontal cortex (dlPFC), few studies have investigated whether these subfunctions are localized to different cortical depths in this region, and none have done so in humans. Here, we use high-resolution functional MRI to interrogate the layer specificity of neural activity during different epochs of a working memory task in dlPFC. We detect activity timecourses that follow the hypothesized patterns: superficial layers are preferentially active during the delay period, while deeper layers are preferentially active during the response. Results demonstrate that layer-specific fMRI can be used in higher-order brain regions to non-invasively map cognitive information processing along cortical circuitry in humans.

## Introduction

Working memory, or the mental capacity to briefly store and manipulate information before acting on it, has been linked to the dorsolateral prefrontal cortex (dlPFC) in both humans and non-human primates (Courtney et al., 1998, 1997, D’Esposito et al., 1995, Goldman-Rakic, 1995). Like much of the cerebral cortex, dlPFC gray matter is organized into layers with distinct cytoarchitecture, connectivity and function. Early electrophysiological work in non-human primates suggested that different epochs of working memory tasks are preferentially associated with activity in different cortical layers (Sawaguchi et al., 1989, 1990). Specifically, it is thought that delay period activity is driven by recurrently connected networks of pyramidal cells in layer III (Goldman-Rakic, 1995), while response-related activity takes place predominantly in layer V (Arnsten et al., 2012, Opris et al., 2011, Wang et al., 2004). Yet direct evidence for this dissociation is limited, due in part to the challenge of separating activity recorded from distinct cortical layers. A recent study in macaques overcame this by using single probes capable of simultaneous recordings from multiple cortical depths (Bastos et al., 2018)

However, to date there is no empirical evidence for such a dissociation in humans, largely because conventional neuroimaging techniques lack the sensitivity and specificity to resolve cortical layers. Recent methodological advances in fMRI, including higher field strengths (i.e., 7 Tesla and above) combined with innovations in pulse sequences and contrast mechanisms, now allow for non-invasive, reliable measurements of cortical depth-dependent activity in humans. These advances have enabled layer-specific imaging in several primary cortices, including visual (Kok et al., 2016, Muckli et al., 2015, Polimeni et al., 2010), auditory (De Martino et al., 2015), and motor (Huber et al., 2017). But it is not yet clear if these techniques are sensitive and robust enough to be applied outside confirmatory studies of primary cortices.

Here, we develop and extend layer fMRI methods to distinguish depth-dependent activity in a region of human association cortex during a cognitive task. Specifically, we use simultaneously acquired blood oxygen level-dependent (BOLD) and cerebral blood volume (CBV) images of dlPFC during working memory. We show that during the delay period, activity is localized to superficial layers, and especially when the task calls for manipulating (as opposed to merely maintaining) information, while during the response period, activity is specifically localized to deeper layers. These results demonstrate the promise of high-resolution fMRI for mapping cognitive cortical circuitry in humans at the mesoscale.

## Results

### Task paradigm

To test our hypotheses about layer-dependent activity in dlPFC, we adapted a well-validated verbal working memory paradigm (D’Esposito et al., 1999). The task contained two types of contrasts (Fig. 1a). In the first contrast (Fig. 1a, top), participants see a string of five random letters (e.g., ‘PXEDL’), then a cue instructing them to either rearrange the letters in alphabetical order (‘ALPHABETIZE’, manipulation condition) or to simply remember them in their original order (‘REMEMBER’, maintenance condition) over the course of a delay period, during which they see only a fixation cross. Finally, a probe letter comes onscreen (e.g, ‘L?’), and participants make a response to indicate the alphabetical or ordinal position of the probed letter. The second contrast (Fig. 1a, bottom) is identical to the first until the response period, at which point participants see either a true probe requiring a button press (e.g., ‘L?’, action condition), or a dummy probe (i.e., ‘∗?’, non-action condition), which indicates that no response is required and they can forget the information associated with that trial. All trials are thus matched for sensory input, with the only difference being the nature of the mental activity during the delay for the first contrast, or the presence or absence of action selection and execution during the response period for the second contrast.

**Fig. 1.**
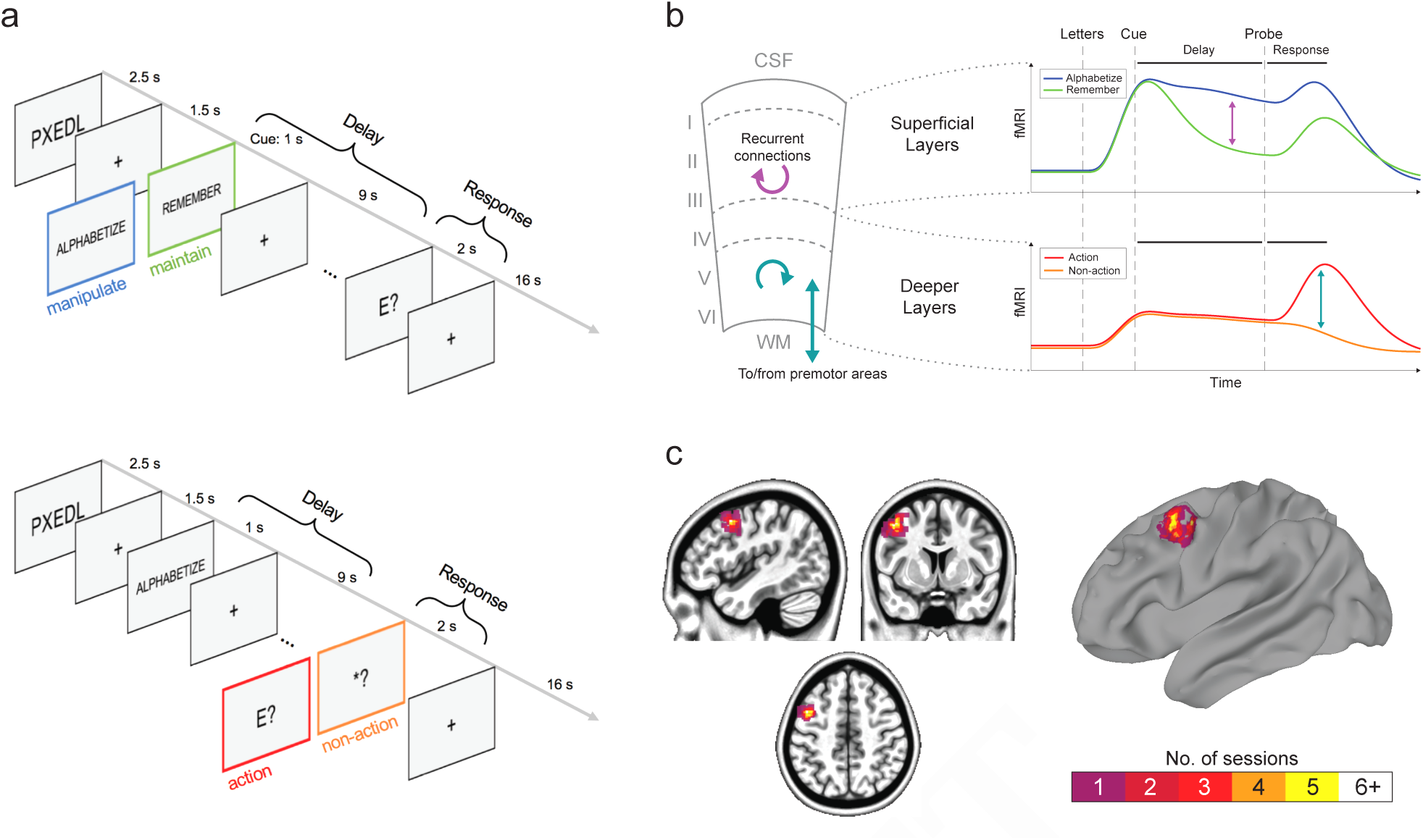
Task, hypothesis and region of interest. (A) Task structure. Top panel: first contrast type, contrasting manipulation (‘alphabetize’) versus maintenance (‘remember’) during the delay period. Bottom panel: second contrast type, contrasting action (true probe) versus non-action (dummy probe) during the response period. Colored frames are for schematic purposes only and were not seen by participants. (B) Schematic of hypothesis: (i) in superficial layers, manipulation trials should evoke more activity than maintenance trials specifically during the delay period due to recurrent excitation in layer III (purple arrows), and (ii) in deeper layers, action trials should evoke more activity than non-action trials due to motor-related functions in layer V (teal arrows). WM, white matter; CSF, cerebro-spinal fluid. (C) Voxelwise overlap of dorsolateral prefrontal cortex regions of interest (ROIs) defined functionally for each individual scan session using the axial readout protocol (total n = 8). ROI overlap is depicted on both volume and surface renderings. The average ROI across participants fell approximately within left Brodmann area 8a.

Thus the task paradigm followed a 2×2×2 design, with trial type (manipulation/maintenance versus action/non-action), period (delay versus response), and cortical depth (superficial versus deep) as the three factors. We hypothesized a triple dissociation between trial type, period, and cortical depth, such that: (1) superficial layers would respond more strongly during the delay period of manipulation trials (as compared to the delay period of maintenance trials), and (2) deeper layers would respond more strongly during the response period of action trials (as compared to the response period of non-action trials). See Fig. 1b for a schematic of the hypothesis. The strength of this experimental design is that we control for each layer’s timecourse of activity primarily by observing the same layer in a different condition, rather than directly comparing activity levels across layers; this helps avoid measurement biases associated with different cortical depths.

### Data acquisition

Nine participants were scanned in a combined total of 13 functional imaging sessions. An additional 14 imaging sessions were conducted for piloting purposes and to collect anatomical data from the same participants (see Methods).

During each high-resolution functional run, we simultaneously measured changes in cerebral blood volume (CBV) and blood-oxygen-level dependent (BOLD) signal using the SS-SI-vascular space occupancy (VASO) method (Lu et al., 2003) with a 3D-EPI readout (Poser et al., 2010) on a 7 Tesla scanner. This method has been implemented to successfully demonstrate layer-specific activity in human motor cortex with good sensitivity and specificity (Huber et al., 2017). The conventional BOLD signal has poor spatial specificity at high resolutions, since it tends to be dominated by large veins at the pial surface. VASO, while it has a lower contrast-to-noise ratio, is a more quantitative measurement that is less biased toward superficial depths. In short, BOLD is more sensitive, while VASO is more specific. Jointly interpreting both contrasts allows for stronger inferences about the magnitude and timing of layer-specific activity than using either on its own (Huber et al.,2018, 2014).

We used two different acquisition protocols over the course of the study. The first had a nominal voxel resolution of 0.9 × 0.9 × 1.1mm (“axial readout protocol”), and was used to quantitatively compare activity timecourses from two distinct cortical depths (superficial versus deep) across participants. Later, we introduced a second, higher resolution acquisition with nominal voxel resolution of 0.76 × 0.76 × 0.99mm (“sagittal readout protocol”) to better visualize activity across different layers in individual participants.

### Location of region of interest

Prefrontal cortex is large, and quite variable across individuals in its structure and functional anatomy. Unlike other cortical landmarks, such as the ‘hand knob’ of the primary motor cortex, functional subdivisions of dlPFC are difficult to pinpoint in individual participants using macroscale anatomical features. Therefore, regions of interest (ROIs) were selected for each participant on the basis of an online functional localizer conducted just prior to the experimental task runs (see Methods).

To better specify our macroscale position within dlPFC, we estimated and visualized the average ROI location across participants (Fig. 1c). Overlap across participants was generally high. The activity of all participants fell within a sphere of 9.5 mm radius (origin at RL 44.2 mm, AP −8.1 mm, IS 47.5 mm), and the peak overlap was centered on a region of the left middle frontal gyrus/superior frontal sulcus corresponding approximately to area 8a, rostral to the frontal eye field (Walker, 1940). In both humans and non-human primates, area 8a is distinguished from areas 8/8b by a more pronounced concentration of large pyramidal cells in layer IIIb, making its cytoarchitecture comparable to neighboring region 9/46d (Petrides and Pandya, 1999), another region classically implicated in WM tasks. It is precisely these pyramidal cells in deep layer III whose recurrent local excitation supports delay-related WM processes (Goldman-Rakic, 1995), making this region a logical place to test our hypotheses.

For each participant, two layers, superficial and deep, were drawn manually within the selected ROI (see Fig. S1 for layer masks for all participants scanned using the axial readout protocol). To ensure that this ROI drawing approach was robust, we conducted test-retest scans separated by several weeks on two participants. Results showed good overlap between ROIs derived from independent experimental sessions (Fig. S4, S5), indicating that the functional region in question can be reliably localized within participants, and the observed layer activity profiles are consistent across sessions. To better specify the position of our “superficial” and “deep” layers with respect to cortical laminae defined cytoarchitectonically, we normalized all available MRI-based anatomical contrasts to an existing histological image (Fig. S3). The boundary between our superficial and deeper layers fell approximately between layer III and layer IV.

### Task performance

Participants performed well on the task (mean accuracy = 77.9 percent, s.d. = 14.4 percent, range = 58.8 – 95.0 percent; note that chance is approximately 20 percent) and there was no difference in accuracy on manipulation versus maintenance trials (paired t-test, *t*_(14)_ = −0.55, p = 0.59). It is therefore unlikely that differences in difficulty between conditions confound the results.

### Activity timecourses

Using data from six participants scanned during eight experimental sessions with the axial readout protocol, we observed layer-dependent activity time-courses that followed the hypothesized patterns: in superficial layers, activity was higher in manipulation relative to maintenance trials during the delay period, and in deeper layers, activity was higher in action versus non-action trials during the response period (Fig. 2). These patterns were visible in both contrasts (BOLD, Fig. 2a; and VASO, Fig. 2b). Below we summarize characteristics of these depth-dependent timecourses during the two main periods of interest, delay and response.

**Fig. 2.**
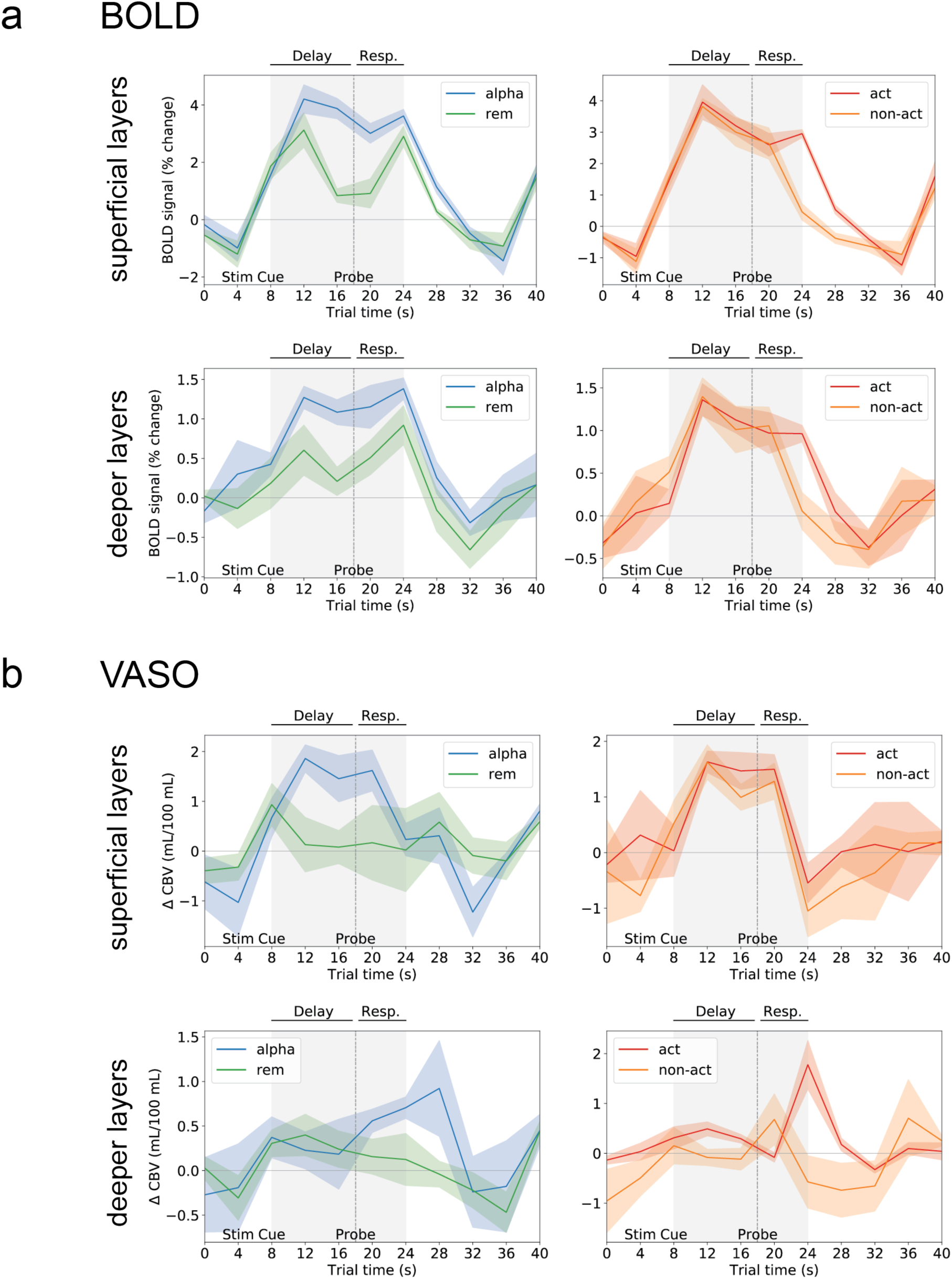
Average activation timecourses for two layers and all four trial types as measured by BOLD and VASO. (A) Left panels show mean BOLD percent signal change in superficial layers (top) and deeper layers (bottom) for the first contrast, which consisted of trial types manipulate (‘alpha’) and maintain (‘rem’). Right panel shows mean BOLD percent signal change in superficial layers (top) and deeper layers (bottom) for the second contrast, which consisted of trial types action (‘act’) and non-action (‘non-act’). (B) Mean activity as measured by VASO (plotsfollowsame schema as in (A)). For both contrasts, lines represent mean and shaded area represents s.e.m. across n = 8 sessions (6 unique participants). Plots are annotated to show timing of trial events: Stim, presentation of letters; Cue, instruction slide (‘alphabetize’ or ‘remember’); Probe, appearance of true letter probe or dummy probe (‘^∗^’).

#### Delay-related activity

In superficial layers, delay-period activity was uniformly high during manipulation trials. This is evident in both BOLD and VASO contrasts, in trials labeled ‘alpha’, ‘action’ and ‘non-action’ (Fig. 2a and 2b, top row; recall that both action and non-action trials call for manipulation, and they are indistinguishable from one another until the probe appears). While it appears from BOLD data as though activity in superficial layers is slightly above baseline even during maintenance trials (Fig. 2a, top left), VASO data indicate little to no response during maintenance trials (apart from a brief initial uptick that may reflect a stimulus-driven sensory encoding signal; Fig. 2b, top left). This agrees with previous reports that human dlPFC is not strictly necessary when the task calls for maintenance only: repetitive transcranial magnetic stimulation (rTMS) to dlPFC selectively impairs manipulation but not maintenance (Postle et al., 2006), and lesions to dlPFC have no effect on maintenance (Mackey et al., 2016).

Compared to superficial layers, deeper layers are markedly less active during the delay. BOLD data appear to indicate some delay-related activity in deeper layers, again more so for manipulation than maintenance trials (Fig. 2a, bottom row). However, VASO data, which have higher spatial specificity, suggest little to no role for deeper layers during this period (Fig. 2b, bottom row). Thus, it seems that delay-related activity occurs predominantly (if not exclusively) in the superficial layers.

#### Response-related activity

During the response period, we observe the opposite pattern: activity in deeper layers is relatively high, but only in trials requiring an action during the response period. Deeper-layer activity peaks at the time of the motor response, which is approximately 6 seconds after the probe comes onscreen (reflecting hemodynamic delay). As expected, this peak is present in action but not non-action trials, and can be seen in both the BOLD (Fig. 2a, bottom right) and VASO (Fig. 2b, bottom right) contrasts.

As for superficial layers, it appears from BOLD data as though their activity remains high through the response period in action trials, while falling off in non-action trials (Fig. 2a, top right). However, this may simply be due to the superficial bias of BOLD, since VASO data (Fig. 2b, top right) indicate that superficial-layer activity is, if anything, suppressed at the response peak in both trial types. This confirms our prediction that response period is preferentially associated with activity in deeper cortical layers.

### Quantification of differential activity

To directly compare activity between trial types, trial periods and cortical depth, we subtracted the average timecourse within each layer between the conditions of interest (i.e., for superficial layers, manipulations¬–maintenance; for deeper layers, action-non-action). Results from both BOLD and VASO confirm that for superficial layers, the difference between manipulation and maintenance peaks during the delay period (Fig. 3a, top and Fig 3b, top), while for deeper layers, the difference between action and non-action trials peaks at the time of the response (Fig. 3a, bottom and Fig. 3b, bottom).

**Fig. 3.**
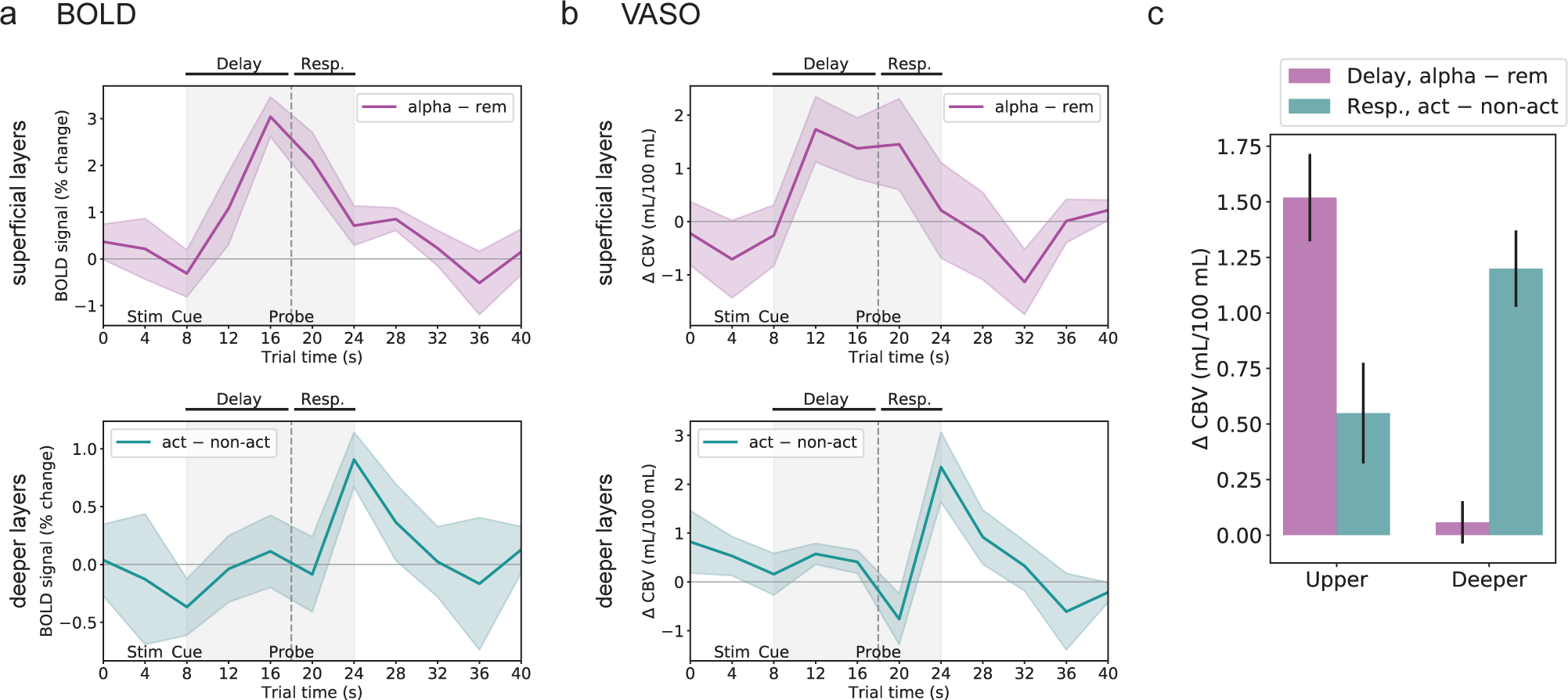
Activation contrasts across layers and conditions of interest. (A) Top: BOLD activity during maintenance (‘rem’) trials subtracted from activity during manipulation (‘alpha’) trials. The largest difference can be seen during the delay period. Bottom: BOLD activity during non-action (‘non-act’) trials subtracted from activity during action (‘act’) trials. The largest difference can be seen during the response period. (B) Subtractions based on VASO activity (plots follow same schema as in (A)). (C) Quantitative comparison of VASO-derived activity levels across layers and trial periods. For all graphs, lines or bars represent mean and shaded area or error bars represents s.e.m. across n = 8 sessions (6 unique participants; same data as in Fig. 2). Plots are annotated as in Fig. 2.

The more quantitative nature of CBV, and the relative lack of depth-dependent biases, permit a direct comparison of VASO-derived activity levels across layers and trial periods. For each cortical depth, we calculated the average differential activity for manipulation over maintenance from measurements acquired during the delay period (timepoints 4, 5 and 6, corresponding to 12, 16 and 20 sec in trial time), and the average differential activity for action over non-action from measurements acquired during the response period (timepoints 7, 8 and 9, corresponding to 24, 28 and 32 sec in trial time). Comparing levels of differential activity (Fig. 3c), it is again evident that superficial layers are more sensitive to the delay-period contrast than the response contrast, while deeper layers are almost exclusively sensitive to the response contrast.

### Visualization of depth-dependent activity

To better visualize the depth-dependent distribution of signal associated with different periods within the trial, we used a second, higher-resolution imaging protocol in which the field of view was a sagittal slab centered on dlPFC with in-plane resolution of 0.76 × 0.76mm. In these experiments, the task consisted exclusively of manipulation/maintenance trials, all requiring an active response (i.e., the first contrast type shown in Fig. 1a, top). Functional signals during manipulation and maintenance trials were investigated across cortical depths.

Layer-dependent activity could be detected in all individual participants imaged using this protocol (n = 5; Fig. 4). Manipulation evoked more activity than maintenance predominantly in superficial layers (green stripes), while signal associated with response (as compared to baseline; red stripes) was predominantly localized to deeper layers. These patterns were visible in both the BOLD (Fig. 4A) and VASO (Fig. 4B) contrasts. Layer ROIs for each participant are shown in Fig. 4C.

**Fig. 4.**
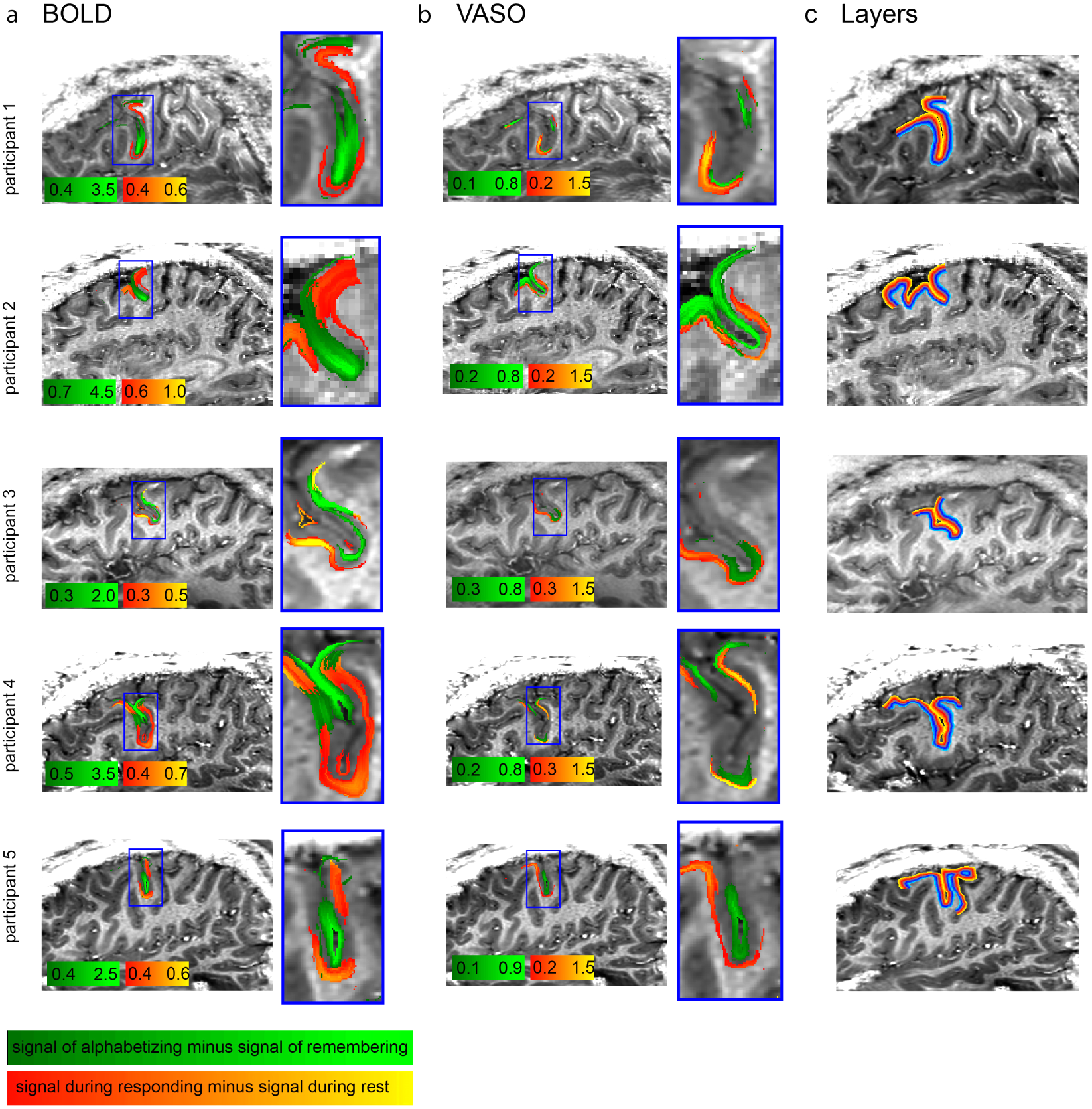
Layer-dependent activation results with sagittal 0.75 mm protocol. Results from five participants using both contrasts, BOLD (A) and VASO (B). Signal changes for delay and response periods are smoothed within layers. No smoothing was applied across layers. Note the different color scales for BOLD and VASO. For an representative layer profile of these maps, see Fig. S3. Estimates of layer (cortical depth) for each participant are shown in (C).

## Discussion

By developing and optimizing state-of-the-art techniques in high-resolution fMRI for cognitive brain areas, we have achieved layer-specific imaging of cortical activity during a working memory task in human dlPFC. We used a three-factor design for which we had clear hypotheses about the location, magnitude and timing of neural activity, and detected timecourses at different cortical depths that followed the expected patterns. Namely, we observed delay-related manipulation activity that was predominantly localized to superficial layers, and response-related activity that was predominantly localized to deeper layers. While working memory has been known to engage dlPFC for decades, the degree to which its subfunctions were layer-specific had been hypothesized but not consistently shown, with few demonstrations even in nonhuman primates (though see Bastos et al. (2018) for recent evidence). Our data interrogate layer-specific functionality directly and non-invasively in humans.

The observed laminar specificity of distinct working memory operations can be interpreted in light of what is known about underlying neural circuitry. First, superficial activity during the delay period likely reflects recurrent excitatory connections. While in early parts of the cortical hierarchy, superficial layers give rise to feedforward connections, at the highest levels (i.e., PFC), laminar projections become more complex. Layer III expands and is the focus of extensive local, recurrent excitatory connections, as well as long-range recurrent connections, e.g., with parietal association cortex (Arnsten et al., 2012, Medalla and Barbas, 2006), which is also heavily involved in working memory. Recurrent excitation among these cells is a feature of their unique molecular profile, notably their preferential expression of n-methyl-d-aspartate (NMDA) receptors and specifically the NR2B subunit, whose slower kinetics allow for persistent firing over long delays; this has been predicted by computational models (Wang, 1999) and confirmed experimentally in primates (Wang et al., 2013).

Second, response-period activity in deeper layers likely reflects functions related to motor control, such as initiating a motor action, suppressing prepotent responses, or a feedback mechanism such as corollary discharge. dlPFC does not project directly to primary motor cortex; rather, it influences motor behavior polysynaptically via higher-order motor areas (Arikuni et al., 1988, Takada et al., 2004). Thus response-related activity in deeper layers may reflect output-circuit activity involving one or multiple such areas. Layer V cells also have dense projections to striatum, which likely also serve to guide movements (Arikuni and Kubota, 1986, Yeterian and Pandya, 1994).

Of note, schizophrenia is associated with altered genetics (reviewed in Arnsten et al. (2012), morphology (Garey et al., 1998, Glantz and Lewis, 2000) and function (Perlstein et al., 2001) in this very dlPFC circuitry. It is hypothesized that decreased delay-related activity in superficial layers, as well as disinhibition in deeper layers, may underlie the deficits in working memory and other cognitive functions seen in these patients. We expect that future studies using layer fMRI in populations with or at risk for schizophrenia will shed new light on the spatiotemporal dynamics of cognitive dysfunction in this illness.

From a methodological perspective, here we used advanced contrast mechanisms and balanced task design to offset differences in vascular physiology across cortical depths, which can introduce substantial biases and limit the interpretability of layer fMRI (Kay et al., 2018). In contrast to gradient-echo BOLD (GE-BOLD), CBV-weighted fMRI signal acquired with VASO allows appropriate separation of microvascular responses at a layer-dependent level (Goense et al., 2012, Kim and Kim, 2010). We avoid biases of different hemodynamic response functions (HRFs) across cortical depths (Petridou and Siero, 2017, Yacoub et al., 2006) by refraining from using statistical general linear model (GLM) deconvolution with predefined HRFs, and by restricting our interpreting to quantitative signal differences that are obtained at the same latency within identical task blocks. Additionally, we collected conventional GE-BOLD fMRI concomitantly with VASO data. The near-simultaneous acquisition of BOLD and VASO data allowed us to obtain a clean BOLD-corrected, CBV-weighed VASO signal. The higher sensitivity of BOLD compared to VASO was helpful in selecting the correct ROI, while the higher spatial specificity of VASO was helpful for interpreting signal across cortical depths.

This work has important implications for non-invasive, *in vivo* mapping of input-output and feedforward-feedback connections in the human neocortex. Outstanding methodological challenges include expanding spatial coverage without sacrificing resolution. Simultaneous imaging of dlPFC, premotor and primary motor cortices would allow for detecting information flow during response generation and execution. Expanding coverage to parietal and sensory areas as well as neighboring prefrontal areas would allow for characterizing interactions that support stimulus perception, information storage and manipulation during the encoding and delay periods. Given current trade-offs between field of view and voxel size, it is still difficult to resolve individual cytoarchitectonic layers (hence we are limited here to drawing conclusions about “superficial” versus “deeper” layers (Fig. S3), rather than the six distinct canonical laminae). But we expect that the ever-advancing tools of high-field fMRI data acquisition and analysis will ultimately transform our understanding of cognition in the awake, behaving human brain.

## Acknowledgements

We thank Amy Arnsten for guidance on experimental design and interpretation. We thank Benedikt Poser and Dimo Ivanov for the 3D-EPI readout that is used in the VASO sequence used here. We thank Andrew Harry Hall and Kenny Chung for administrative support of human volunteer scanning. We thank Sriranga Kashyap for helpful tips on adjusting manual initial registration used to generate Fig. 1c and Fig. S4-5. Portions of this study used the high-performance computational capabilities of the Biowulf Linux cluster at the National Institutes of Health, Bethesda, MD (biowulf.nih.gov).

## Authors’ contributions

E.S.F. conceptualized the study. E.S.F., L.H., and D.C.J. designed the experimental paradigm. L.H. designed and optimized the data acquisition and analysis methodology. L.H. and E.S.F. collected the data and performed the analyses. E.S.F., L.H., and D.C.J. generated data visualizations. P.A.B. supervised study design and interpretation. E.S.F. wrote the original draft, with contributions from L.H. and revisions from D.C.J. and P.A.B.

## Data and materials availability

Data and code will be made available online at publication <*link to be inserted here*>.

This preprint is formatted based on a L^A^T_E_Xclass by Ricardo Henriques.

## Supplementary Material

### Methods

#### Participants

Nine healthy volunteers (age 20-47 years at the time of the experiment) participated after granting informed consent under an NIH Combined Neuroscience Institutional Review Board-approved protocol (93-M-0170, ClinicalTrials.gov identifier: NCT00001360) in accordance with the Belmont Report and US federal regulations that protect human subjects. Three participants were male and six were non-pregnant females.

The functional part of this study consisted of 34 hours of scan time from 17 two-hour scan sessions. Due to the inter-participant anatomical variability in the sub-millimeter mesoscale, the scan slots were used to do multiple comprehensive experiments in the same participants, rather than short experiments in a larger cohorts. This is consistent with previous layer fMRI studies (De Martino et al., 2015, Goense et al., 2012, Huber et al., 2017, Kim and Kim, 2010, Kok et al., 2016, Muckli et al., 2015). Four scan sessions were used as pilot experiments to optimize the task design and investigate motion limitations and sequence artifact level. In the remaining functional sessions, nine volunteers participated in a combined total of 13 functional imaging sessions. Four of the participants were invited multiple times (for up to three imaging sessions) to confirm the reproducibility of the results (activation location). All nine participants were invited for a separate scan session to obtain high-resolution reference data at 0.7 mm T1-weighted images with an MP2RAGE sequence (Marques et al., 2010). The four volunteers that participated in multiple functional sessions were also re-invited for an ultra-high-resolution T1 and T2^∗^ reference scan with 3-7 averages of 0.5 mm and 0.4 mm resolution, respectively. This totals to data from 62 hours scan time.

#### Task paradigm

The task was created using PsychoPy software (Peirce, 2007). For the axial readout protocol (TR = 2 s, described below), each trial consisted of the following epochs (example, duration): letter string presentation (PXEDL, 2.5 s), fixation cross (+, 1.5 s), instruction cue (ALPHABETIZE or REMEMBER, 1 s), delay period with fixation cross (+, 9 s), probe (D? or ^∗^?, 2 s), inter-trial interval with fixation cross (+, 16 s). Participants could register a response at any time following the appearance of the probe and before the start of the next trial (i.e., anytime during the inter-trial interval). Each trial thus lasted 32 s, and each run consisted of 20 trials plus brief (8 s) additional fixations at the beginning and end of the run, for a total of 10:56 min:sec per run. Runs alternated between two contrast types: (1) manipulation versus maintenance (consisting of a mix of ALPHABETIZE and REMEMBER trials, all requiring action), and (2) action versus non-action (consisting of a mix of action and non-action trials, all ALPHABETIZE). Within each run, the 10 trials of each type were presented in a pseudorandom order that was the same for all runs, to facilitate averaging.

For the higher-resolution sagittal readout protocol (described below), trial epoch timings were adjusted to match the longer TR (2.5 s) by scaling the duration of each epoch by a multiplier of 1.25. Each trial thus lasted 40 s, and the duration of these runs was 13:40 min:sec. All other parameters, including the pseudorandom order, were kept the same as above.

Prior to the start of the experimental runs, we ran a 6-minute functional localizer consisting entirely of ALPHABETIZE trials and slightly altered timing. The length of all trial epochs was as described above except the inter-trial interval, which was shortened to 5 s to create a 10-s on, 10-s off paradigm. Delay-related activity (including cue plus delay-related fixation) was considered signal, while all other trial epochs were treated as baseline. This initial 6-minute experiment was conducted at standard resolution and analyzed in real time, allowing us to functionally define a region of interest within dlPFC in each individual participant while the participant was in the scanner. The location of peak activity from the real-time analysis was used to position the coverage of the subsequent sub-millimeter experiments.

#### Experimental setup

All imaging was performed on a MAGNETOM 7T scanner (Siemens Healthineers, Erlangen, Germany) with a single-channel-transmit/32-channel-receive heal coil (Nova Medical, Wilmington, MA, USA). Imaging sessions did not exceed 120 minutes. Imaging slice position and slice angle were adjusted individually for every participant on the basis of the functional localizer described above. A 3rd-order *B*_0_-shim was done with three iterations using vendor-provided tools. The shim volume covered the entire imaging field of view (FOV) and was extended down to the circle of Willis in order to obtain sufficient *B*_0_-homogeneity to exceed the adiabaticity threshold of the inversion pulse.

Following the functional localizer, run type alternated between the first contrast (manipulation/maintenance) and the second contrast (action/non-action). Most participants completed five runs (3 of the first contrast and 2 of the second); when time allowed, a sixth run was acquired (second contrast).

#### Axial readout protocol

Six unique participants were scanned during eight sessions using this protocol (including test-retest for two participants). In-plane voxel resolution was 0.9 mm with a slice thickness of 1.1 mm. The protocol parameters were as follows: Readout type: 3D-EPI with one segment per k-space plane (Poser et al., 2010), in-plane resolution 0.91 × 0.91mm^2^, slice thickness 1.1 mm, FLASH GRAPPA 3, partial Fourier in the first phase encoding direction: 6/8, no partial Fourier in the second phase encoding direction, TR_*V ASO*_ = 2000 ms, TR_*V ASO+BOLD*_ = 4000 ms, FOV read and phase =150 mm, matrix size = 162, TE = 20 ms, read bandwidth = 1144 Hz/Px, phase echo spacing = 0.98. Assuming a gray-matter T2^∗^ = 28 ms, the expected T2^∗^ blurring for EPI-readout results in a signal leakage of 12 percent from one voxel into the neighboring voxels along the first phase-encoding direction. A more detailed list of scan parameters can be found on GitHub: https://github.com/layerfMRI/Sequence_Github/blob/master/DLPFC_Emily/Emily_Intermediate_protocol.pdf

#### Sagittal readout protocol

Five unique participants were scanned using this protocol. The protocol parameters are as follows: Readout type: 3D-EPI with one segment per k-space plane (Poser et al., 2010), in-plane resolution 0.75 × 0.75*mm*^2^, slice thickness 0.99 mm, FLASH GRAPPA 3, partial Fourier in the first phase encoding direction: 6/8, no partial Fourier in the second phase encoding direction, *TR_V_ ASO* = 2500 ms, *TR_V ASO+BOLD_* = 5000 ms, FOV read = 130 mm, FOV phase 98.8 percent, matrix size = 172, TE = 27 ms, read bandwidth = 908 Hz/Px, phase echo spacing = 1.23 (limited by peripheral nerve stimulation thresholds). Assuming a gray-matter T2^∗^ = 28 ms, the expected T2^∗^ blurring for EPI-readout results in a signal leakage of 14 percent from one voxel into the neighboring voxels along the first phase-encoding direction. A more detailed list of scan parameters used can be found on GitHub: https://github.com/layerfMRI/Sequence_Github/blob/master/DLPFC_Emily/DLPFC_high_res_076_0.76_1.pdf

#### VASO-specific protocol parameters

Both readout protocols were acquired with the same VASO preparation module. The protocol parameters were: Inversion pulse type: TR-FOCI pulse with a bandwidth of 6.4 kHz, *μ*, = 7, pulse duration: 10 ms, non-selective. The phase skip of the adiabatic inversion pulse was adjusted to 30 deg to achieve an inversion efficiency of 80 percent, shorter than the arterial arrival time in the dlPFC (Mildner et al., 2014). The inversion time was adjusted to match the blood-nulling time of 1100 ms as done in previous studies (Huber et al., 2017). To account for the T1-decay during the 3D-EPI readout and potential related blurring along the segment direction, a variable flip angle was chosen. The flip angle of the first segment was adjusted to be 22 deg. The subsequent flip angles where exponentially increasing, until last k-space segment was excited with a desired flip angle of 90 deg.

#### Image reconstruction

Image reconstruction was done in the vendor-provided platform as done previously (Huber et al., 2017). GRAPPA 3 kernel fitting was done on FLASH ACS data, using a 3 × 4 kernel, 48 reference lines, and regularization parameter *χ* = 0.001. RF-channels were combined with the sum-of-squares. To minimize resolutions losses in the phase-encoding direction due to T2^∗^-decay partial, Fourier reconstruction was done with POCS using 8 iterations.

#### Anatomical reference data

In separate scan sessions, 0.7 mm resolution T1-maps were collected covering the entire brain with an MP2RAGE sequence (Marques et al., 2010) for every participant. These data were not used in the functional pipelines to delineate ROIs for assessing layer-dependent activity changes. Instead, these images were used to investigate the reproducibility of location of activity across sessions (Figs. S4, S5) and across participants (Fig. 1c).

In four of the participants that were invited for more than two 2-hour sessions, slab-selective isotropic 0.5 mm and 0.4 mm resolution anatomical data were collected with MP2RAGE and Multi-Echo FLASH, respectively. Those anatomical data were not used in the pipeline for generating cortical profiles. They are used to compare and validate the approximate position of the cyto-architectonically defined cortical layers of individual participants to the 20 reconstructed cortical depths in which the functional data are processed (Fig. S3).

#### Functional image preprocessing

For a schematic overview of the analysis pipeline, see Fig. S2. DICOM images were converted to NIFTI using the ISISCONV converter (Fig. S2a). Motion correction was performed using SPM software (Statistical Parametric Mapping; SPM12, Friston et al. (1994)) and was done separately for nulled and not-nulled frames (Fig. S2b). A 4th order spline function was used for spatial interpolation. Motion correction and registration across runs was done simultaneously. This minimized the effect of spatial resolution loss to one single resampling step (Polimeni et al., 2018). Motion traces of nulled and not-nulled were visually inspected to ensure good overlap for the two contrasts (Fig. S2b). Data from one participant were excluded at this step due to motion exceeding 5mm, leaving a total of n = 8 experimental sessions from 6 unique participants for timecourse analysis.

#### Timecourse extraction

Following preprocessing, frames were sorted into their respective contrast: not-nulled (BOLD) or nulled (VASO; Fig. S2c). Next, runs of the same contrast type were averaged (Fig. S2d), and within these average runs, trials of the same type were averaged (Fig. S2e). Because all runs have the same trial order, and all trials have the same epoch structure and timing, runs and trials can be averaged without deconvolving the hemodynamic response. This is an important feature of our experimental design, since hemodynamic responses differ across cortical depths (Petridou and Siero, 2017). Following trial averaging, VASO data were BOLD corrected using the dynamic division method (Fig. S2e). Thus, for each contrast (BOLD and VASO), each participant had four average trials: alphabetize, remember, action, and non-action.

In a parallel analysis, a region of interest (ROI) in the left dlPFC was defined for each participant (Fig. S2f). The approximate location of the ROI was taken from the 6-minute functional localizer (Fig. S2f, left) following GLM analysis with FSL FEAT (Version 5.98; Worsley (2001)). For the complete FEAT design protocol, please see (https://github.com/layerfMRI/repository/tree/master/DLPFC_Emily/Featdesign). The ROI was manually selected and drawn for every individual participant (see Fig. S1 for drawn ROIs in every participant). Rather than acquire an additional T1-weighted image for anatomical reference, we used the functional EPI data itself to estimate the T1 contrast, and used this for manual delineation of two layers within this ROI, one superficial and one deep (Fig. S2f, right). The advantage of this approach is that it avoids the distortion correction and resampling steps necessary for registering EPI images to a separately acquired T1 image, preserving spatial specificity.

Next, at each timepoint, signal was averaged across all voxels within each layer—superficial and deeper—to derive one average timecourse per layer in each of the four trial types. Thus, each participant had eight timecourses: one per layer (upper, deeper) per trial type (alphabetize, remember, action, non-action; Fig. S2g).

#### Timecourse normalization

Before pooling data across subjects, BOLD timecourses were normalized within subjects using the following steps. First, a per-participant mean active BOLD signal (*m_a_*) was calculated by averaging all active timepoints across all eight layer-condition timecourses (where “active” refers to the 8 timepoints beginning with letter presentation and ending 10 seconds after the appearance of the probe, the point at which signal is expected to have returned to at or near baseline). Next, the BOLD signal *b* at each timepoint *t* was normalized as follows: 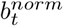 = (*b_t_* – l)/(*m_a_* – 1). This step serves to center BOLD activity approximately around zero, such that active and baseline timepoints have positive and negative values, respectively.

Following this within-subject normalization, the mean and standard deviation of each time point were calculated for each of the eight layer-condition combinations across participants. From these average timecourses, a grand mean was calculated for active timepoints (*M_a_*) as well as baseline time-points (*M_b_*, defined as all non-active timepoints, or the first timepoint prior to letter presentation plus the penultimate and ultimate timepoints of each trial). This baseline value *M_b_* was then subtracted from all values, and the resulting values were multiplied by (*M_a_* – 1) ∗ 100 to yield values interpretable as percent signal change. Standard error was calculated as the standard deviation across participants divided by 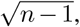, where n = 8, reflecting the number of experimental sessions contributing data points.

Note that unlike BOLD, VASO is a quantitative measure that is proportional to a physical unit (mL per 100 mL tissue volume). Thus, it is not necessary to perform within-subject normalization based on active timepoints before pooling data for group analysis. VASO data were instead transformed as follows. For each participant, the VASO signal v at each time-point t was normalized as: 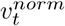 = –*v_t_* ∗ 100. Following this step, VASO signal values were averaged across participants to derive eight average timecourses. Because VASO is a negative contrast (such that a decrease in signal reflects greater neural activity), these signal values were multiplied by −1 to facilitate interpretation, and the mean baseline signal was subtracted from all values to yield the final set of time-courses. As above, standard error was calculated as the standard deviation across participants divided by 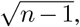, where n = 8.

Note that the functional contrast used here to compare activity in superficial and deeper layer corresponds to normalized signal difference between conditions. No inferential statistical thresholds were used at any point along the functional signal analysis. We refrained from using inferential statistical models to measure activation, to avoid biases of variable noise magnitudes and hemodynamic response function timings across cortical depths.

#### Layering for sagittal protocol

Cortical depths were estimated directly in EPI space without alignment to so-called anatomical space. This minimizes the risk of resolution loss due to multiple spatial resampling steps and avoids any potential errors in distortion correction and registration. An anatomical reference contrast was calculated from the functional data by calculating the inverse signal variability across nulled and not-nulled images, divided by the mean signal. This measure is called here T1-EPI and provides a good contrast between white matter (WM), gray matter (GM) and cerebro-spinal fluid (CSF; see background images in Figure 4, S1 and S3). Borderlines between GM/WM and GM/CSF are manually drawn based on this contrast. Manually drawn border lines are shown for all participants in Figure 4c (bright yellow for GM/CSF and bright blue for GM/WM). Twenty-one layers were calculated between these borderlines with the LAYNII program LN_GROW_LAYERS: https://github.com/layerfMRI/LAYNII. In order to minimize partial volume effects and allow the calculation of smooth layers, the layering calculation was applied on a fourfold finer grid that the native functional resolution. This means that the number of layers is higher than the number of independent voxels sampled across the cortical depth. The number of layers should not be confused with the effective resolution across cortical depths. Given the cortical thickness of 3.5-4 mm in dlPFC (Fischl and Dale, 2000, Williams and Goldman-Rakic, 1993), the resolution of 0.76 mm inplane and 0.99 mm slice thickness, is sufficient to sample 3-6 independent voxels across cortical depth. This is enough to estimate activity in superficial and deeper layers (red-yellow compared to blue-turquoise in Fig. 4c) with Nyquist sampling. For best visibility, fMRI signals were smoothed along the tangential direction of the cortex with a Gaussian kernel of 0.76 mm. In order to maintain the spatial specificity across layers, no smoothing was applied across cortical depths. The functional results of this layer-analysis are shown in Fig. 4.

#### Spatial alignment across participants

To investigate the consistency of the location of activity across days and across participants, the layer masks and the corresponding activation maps were transformed into anatomical reference spaces. Registration was done with SyN in ANTs (Advanced Normalization Tools; Avants et al. (2008)) with a spline interpolation. Since the imaging coverage of the functional data is significantly smaller than the whole brain, it was necessary to provide a manual starting point for the ANTs registration to converge on reasonable registration quality. The initial manual registration was done in ITK-SNAP. The registration from EPI-space to the participant specific anatomical space was done by means of the similar T1 contrast of T1-EPI and the MP2RAGE UNI-DEN image. The same spatial operation was applied to the layer masks and the functional activation maps. The resulting activation patterns were compared across days in the anatomical space of individual participants (Figs. S4, S5).

For comparisons of activation locations across participants, the binary layer masks where further registered to MNI-template space. The participants’ MP2RAGE UNI-DEN contrast images were registered to the MNI1 mm template brain provided in FSL. This was done in ANTs using an affine registration. The same affine transformation was applied to the layer masks using a nearest neighbor interpolation. The overlap across participants of the resulting binary masks was computed and visualized to estimate the spatial consistency (Fig. 1c).

**Fig. S1.**
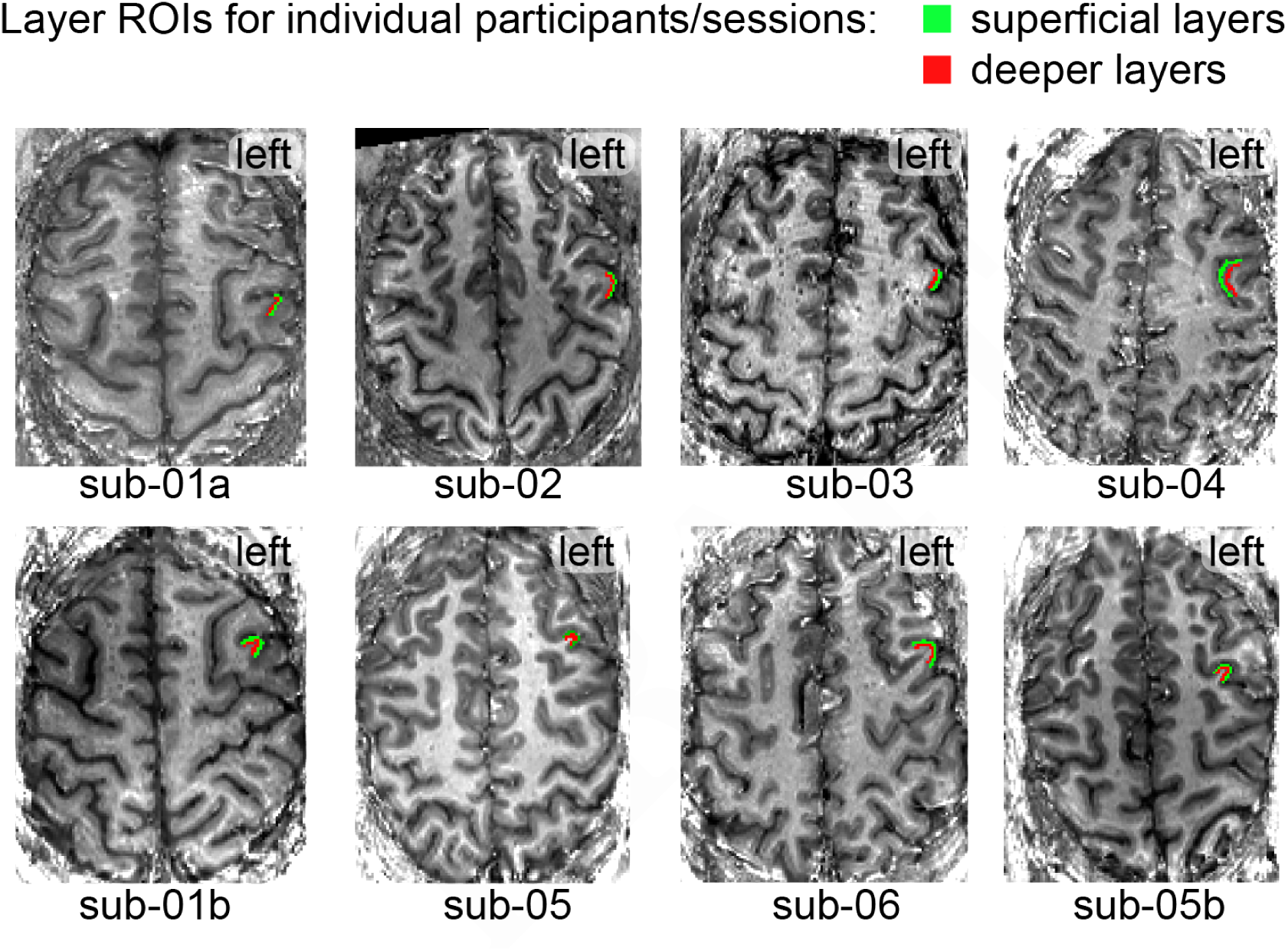
Layer ROIs for individual participants and sessions. Masks of superficial and deeper layers for each participant/session that were used to extract the time courses in Fig. 2-3. The grayscale background contrast refers to the T1-contrast in functional VASO data. Here the T1-contrast in the functional data is used to identify superficial and deeper voxels (manually drawn green and red masks).

**Fig. S2.**
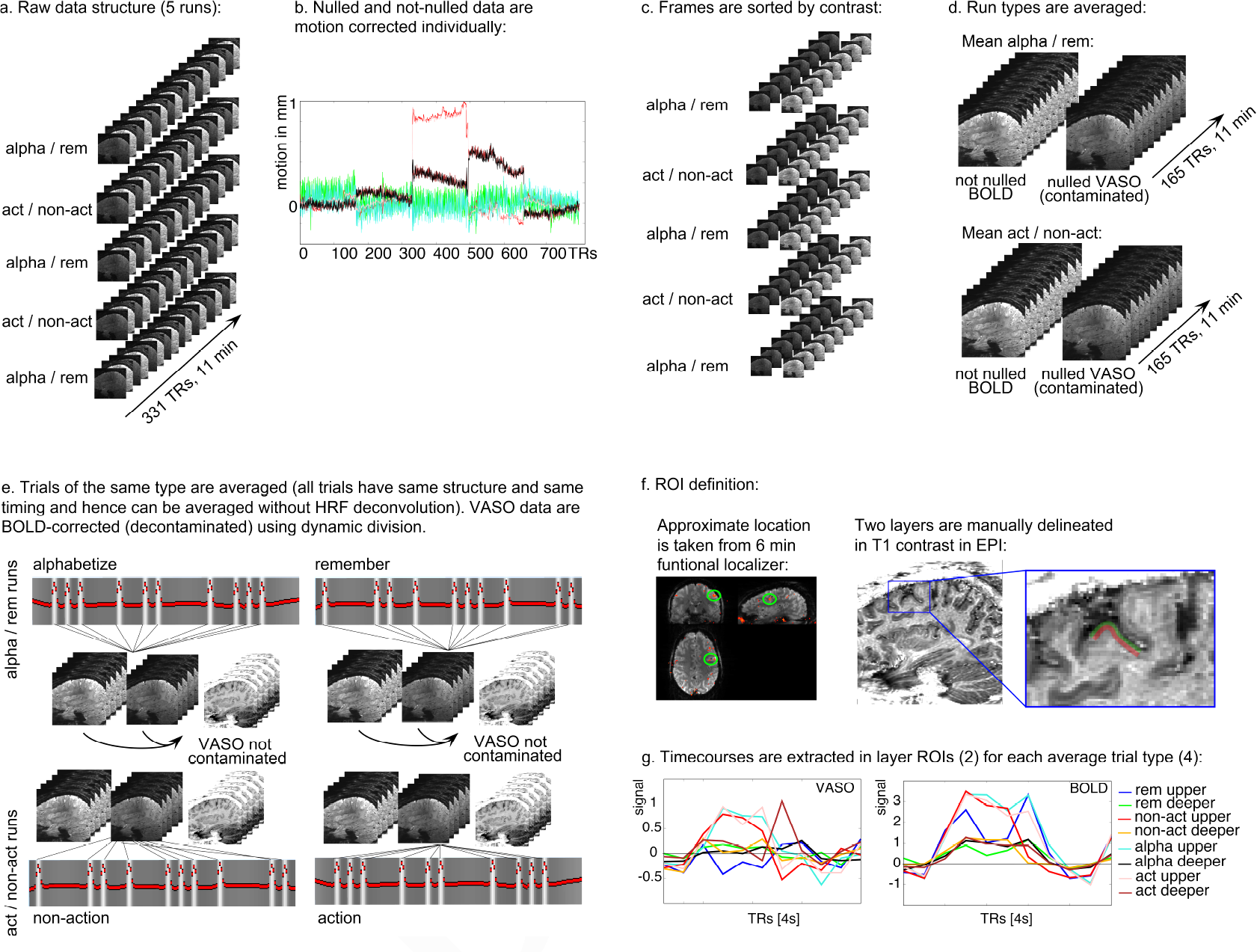
Graphical depiction of the analysis pipeline. A) Multiple runs are collected per session with two different tasks. Each run consists of interleaved images with and without blood nulling. B) Nulled and not-nulled images are motion corrected separately for the whole session. It is manually checked that the motion traces of nulled and not-nulled images are matching. C) The time series are sorted by imaging contrast. D) Runs of the same type are averaged within contrast. E) Time series from all trials are averaged based on their task condition. In the 2 × 2 used here, this results in 4 average trial types. Nulled images are corrected for BOLD contamination with a time-wise division of not-nulled images to provide a clean VASO contrast. F) Based on the approximate location of activation in the low-resolution functional localizer, two layer ROIs are manually drawn on the T1-EPI anatomical images. G) Time series for all four task conditions are extracted from the ROI of superficial and deeper layers for BOLD and VASO contrasts.

**Fig. S3.**
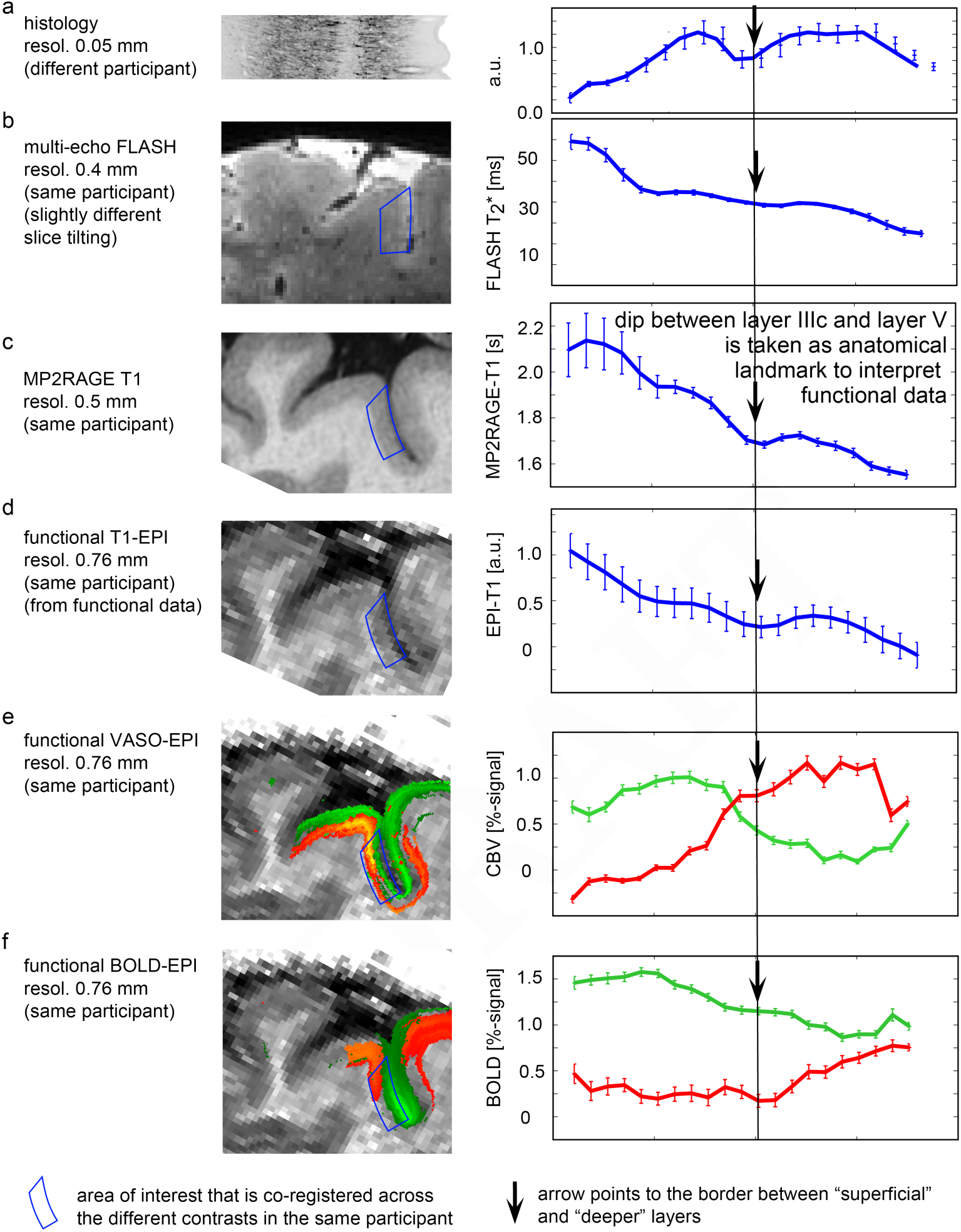
Normalization of cortical depth to cytoarchitectonic cortical layers. The position of the transition area between layer III and IV in a histological section of Brodmann area 8 can be identified with a local dip (black arrow in A). This landmark is also visible in in-vivo T1 and T2^∗^ weighted profiles (B-D). It particularly pronounced in 0.5 mm T1-profiles from MP2RAGE, a contrast that can be compared to the inverse VASO weighting of the functional data (D). This dip of cytoarchitectonic layer III and IV is used here as the border between so-called “superficial layers” and “deeper layers” which show different responses to the different functional task contrasts (E-F).

**Fig. S4.**
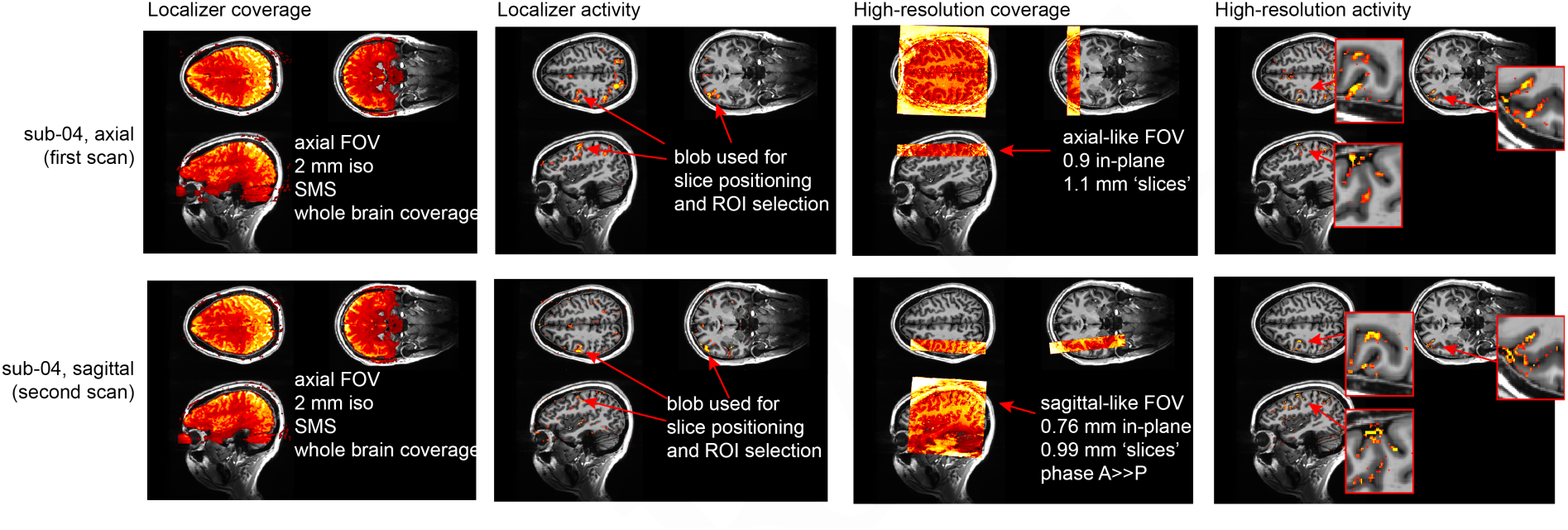
Reproducibility of activation loci in one participant across the axial and sagittal imaging protocols acquired on two different days. It can be seen that the same parts of gray matter are engaged in the task across the two different acquisition protocols (axial-like, top row; and sagittal-like, bottom row) acquired on two different days. The consistency of the locus of activation is within the accuracy of the registration quality.

**Fig. S5.**
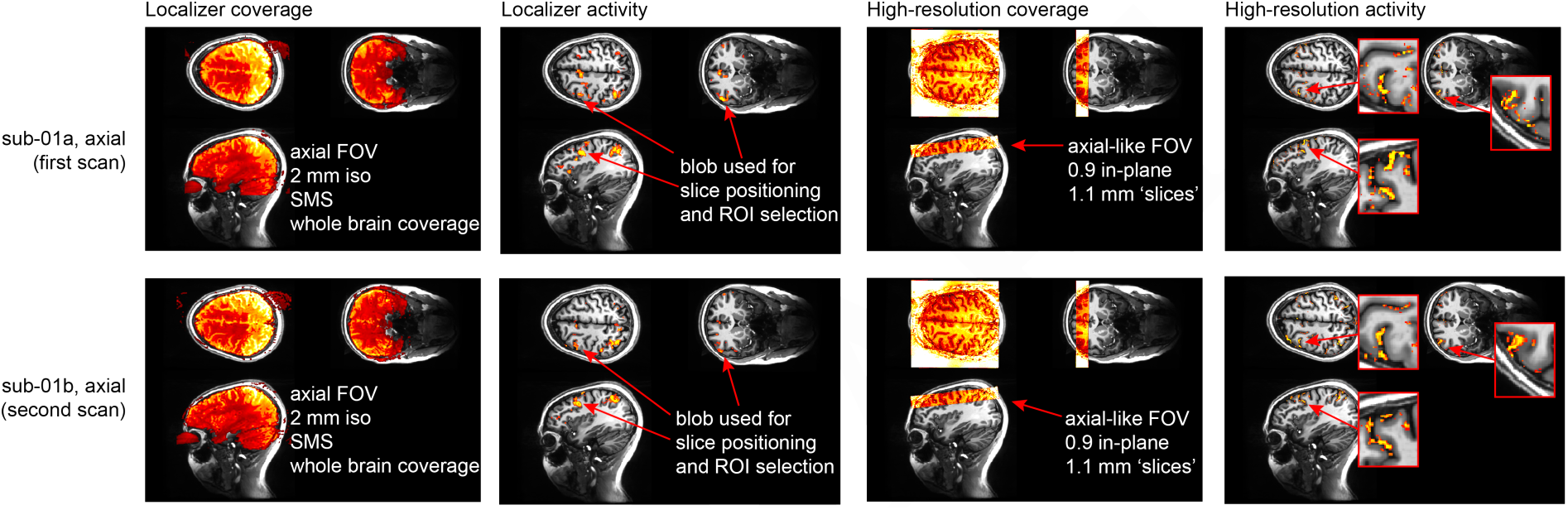
Reproducibility of activation loci in one participant scanned with the axial imaging protocol on two different days. It can be seen that the same parts of gray matter are engaged in the task across the same acquisition protocol on two different days (first session, top row; second session, bottom row). The consistency of the locus of activation is within the accuracy of the registration quality.

